# Skeletal Muscle mTORC1 Activation Increases Energy Expenditure and Reduces Longevity in Mice

**DOI:** 10.1101/720540

**Authors:** Erin J. Stephenson, JeAnna R. Redd, Detrick Snyder, Quynh T. Tran, Binbin Lu, Matthew J. Peloquin, Molly C. Mulcahy, Innocence Harvey, Kaleigh Fisher, Joan C. Han, Nathan Qi, Alan R. Saltiel, Dave Bridges

## Abstract

The mechanistic target of rapamycin (mTORC1) is a nutrient responsive protein kinase complex that helps co-ordinate anabolic processes across all tissues. There is evidence that signaling through mTORC1 in skeletal muscle may be a determinant of energy expenditure and aging and therefore components downstream of mTORC1 signaling may be potential targets for treating obesity and age-associated metabolic disease. Here, we generated mice with *Ckmm-Cre* driven ablation of *Tsc1*, which confers constitutive activation of mTORC1 in skeletal muscle and performed unbiased transcriptional analyses to identify pathways and candidate genes that may explain how skeletal muscle mTORC1 activity regulates energy balance and aging. Activation of skeletal muscle mTORC1 produced a striking resistance to diet-and age-induced obesity without inducing systemic insulin resistance. We found that increases in energy expenditure following a high fat diet were mTORC1-dependent and that elevated energy expenditure caused by ablation of *Tsc1* coincided with the upregulation of skeletal muscle-specific thermogenic mechanisms that involve sarcolipin-driven futile cycling of Ca^2+^ through SERCA2. Additionally, we report that constitutive activation of mTORC1 in skeletal muscle reduces lifespan. These findings support the hypothesis that activation of mTORC1 and its downstream targets, specifically in skeletal muscle, may play a role in nutrient-dependent thermogenesis and aging.

## Introduction

Obesity is a worldwide health problem, with comorbidities including diabetes, cardiovascular and liver disease [1]. Current modalities to prevent or reverse obesity are ineffective and short-lived, either due to poor adherence to lifestyle interventions or reductions in energy expenditure and increases in hunger after weight loss [2–4]. The genetic and dietary modifiers of energy expenditure are not well understood, but there is evidence that signaling through the mechanistic Target of Rapamycin Complex 1 (mTORC1) may play a role [5–8].

mTORC1 is a nutrient responsive protein kinase complex expressed in all known eukaryotic cells. This complex is activated by anabolic signals including insulin, amino acids and energy abundance, and repressed during periods of energy and nutrient deprivation (see [9] for review). mTORC1 integrates these signals, and helps co-ordinate anabolic processes such as protein synthesis, lipogenesis [10–12], glycogenesis [13] and cellular differentiation [14–16], while also promoting insulin resistance [10,17]. These effects are often tissue-specific, reflecting the cell-type specific responses to elevated nutrient and energy status.

Skeletal muscle is the major site of postprandial glucose disposal and the primary determinant of resting energy expenditure in mammals [18,19]. Constitutive activation of mTORC1, via muscle-specific deletion of its negative regulator *Tsc1*, results in age-related myoatrophy, dysregulation of autophagy induction and increased expression of mitochondrial enzymes [6,20,21]. Consistent with the latter, cell culture models implicate mTORC1 as a positive regulator of mitochondrial biogenesis and aerobic ATP production [22–24]. During the aging process, skeletal muscle exhibits a fiber-type transformation towards a more oxidative phenotype, concomitant with increased mTORC1 activity. In line with these observations, several studies have implicated mTORC1 inhibition as a mechanism of organismal lifespan extension in yeast, worms and mammals [25–27]; however, the tissue or tissues that link mTORC1 activity to lifespan have not yet been identified.

Skeletal muscle is an important tissue for understanding aging, insulin sensitivity and changes in energy metabolism, as functional differences in muscle strength predict lifespan in humans [28–33]. Furthermore, mTORC1 regulates several important metabolic processes in muscle; including oxidative stress, the unfolded protein response, autophagy and lipid metabolism [20,34,35]. Here, we have performed unbiased transcriptional analyses to identify pathways and candidate genes that may explain how skeletal muscle mTORC1 activity regulates energy balance and aging. We show that chronic mTORC1 activation in skeletal muscle (via deletion of its negative regulator, *Tsc1*) promotes increased energy expenditure, but reduced lifespan.

## Materials and Methods

### Animals

All mice were purchased from The Jackson Laboratory. Unless otherwise stated, animals were fed a normal chow diet from Harlan Teklad (catalog # 7912). For high fat diet (HFD) studies, animals were provided *ad libitum* access to a diet with 45% of calories from fat (Research Diets D1492). HFD feeding was initiated when animals were approximately 10 weeks of age. For tissue collection, animals were anesthetized with isoflurane before being sacrificed by cervical dislocation at 25 weeks of age.

Muscle-specific *Tsc1* knockouts were generated by crossing FVB-Tg(*Ckmm*-*Cre*)5Khn/J transgenic mice (stock 006405) with floxed *Tsc1*^*tm1Djk*^/J mice (stock 005680). To generate F1 mice that were heterozygous for the floxed allele, mice that either possessed or lacked the *Ckmm-Cre* transgene were intercrossed to generate knockout mice (*Tsc1*^*fl*/fl^, *Ckmm-Cre*^Tg/+^), wild type mice (*Tsc1*^*+/+*^, *Ckmm-Cre*^+/+^) and controls containing the transgene only (*Tsc1*^*+/+*^, *Ckmm-Cre*^Tg/+^), or the floxed allele only (*Tsc1*^*fl*/fl^, *Ckmm-Cre*^+/+^). All four genotypes were studied in all experiments. If there were no significant differences between the three control genotypes, these results were combined and labeled as wild-type. Animals were sacrificed in either the fed or fasted state as indicated in the figure legends, at approximately ZT3. All animal procedures were carried out in accordance with the National Institute of Health guide for the care and use of Laboratory animals and were approved by The University of Michigan and UTHSC Institutional Animal Care and Use Committees prior to the work being performed.

### Body Composition and Indirect Calorimetry

Body weights were determined using a standard scale, whereas body composition was determined in conscious animals by magnetic resonance (EchoMRI 1100, EchoMRI, Houston, TX). Adipose tissue weights (dorsolumbar-inguinal and gonadal depots) were dissected from both the left and right sides (the combined weight of both sides is reported). For indirect calorimetry studies, physical activity, VO_2_ and VCO_2_ were determined using a home-cage style Comprehensive Laboratory Animal Monitoring System (CLAMS, Columbus Instruments, Columbus, OH) with hanging feeders containing pelleted food, under light and temperature-controlled conditions (12:12hr, 25ºC). Ambulatory activity was calculated as the sum of x and y axis beam breaks. The first 6h of CLAMS measurements were discarded to accommodate acclimation, after which continuous measurements were made over three consecutive days. Data were analyzed by mixed linear models with considerations [36,37]. To account for the dominant effect of lean mass on total energy expenditure, lean mass was included as a covariate in the mixed linear models. Energy expenditure was calculated as heat, using the Lusk equation in Oxymax software (Columbus Instruments, Columbus, OH), and rates of carbohydrate and lipid oxidation were calculated according to Péronnet and Massicotte [38] with the assumption that the rate of protein oxidation occurring under standard housing conditions is negligible. For rapamycin treatments, animals were individually housed for 10 consecutive days (days 3-10 were in CLAMS cages). Mice received four days of vehicle treatment (1% Tween, 1% PEG-8000), followed by three days of treatment with either vehicle or the selective mTOR inhibitor rapamycin (3 mg/kg/d, via intraperitoneal injection). Mice were then switched to HFD and indirect calorimetry measurements continued for an additional three days, during which time mice continued to receive daily injections of either vehicle or rapamycin.

### Hyperinsulinemic Euglycemic Clamp and Tissue 2-Deoxyglucose Uptake

Animals were anesthetized with sodium pentobarbital (50−60 mg/kg, given intraperitoneally) and indwelling catheters were implanted into the right jugular vein and the right carotid artery. The catheters were tunneled subcutaneously and exteriorized at the back of the neck via a stainless-steel tubing connector that was fixed subcutaneously upon closure of the incision. Animals were allowed to recover and mice with healthy appearance, normal activity, and weight regain to or above 90% of their pre-surgery levels were used for the study. Experiments were carried out in conscious and unrestrained animals using techniques described previously [39–42]. The primed (1.0 µCi)-continuous infusion (0.05 µCi/min and increased to 0.1 µCi/min at t = 0) of [3-^3^H] glucose (50 µCi/ml in saline) was started at t = −120min. After a 5-6 hour fast, the insulin clamp was initiated at t = 0, with a prime-continuous infusion (40 mU/kg bolus, followed by 2.5 mU/kg/min) of human insulin (Novo Nordisk). Euglycemia (120~130 mg/dL) was maintained during the clamp by measuring blood glucose every 10 min and infusing 50% glucose at variable rates, accordingly. Blood samples were collected from the right carotid artery at t = 80, 90, 100, and 120 min for determination of glucose specific activity. Blood insulin concentrations were determined from samples taken at t = −10 and 120 min. A bolus injection of [1-^14^C]-2-deoxyglucose ([^14^C]2DG; PerkinElmer) (10 µCi) was given at t = 120 min. Blood samples were taken at 2, 5, 10, 15, and 25 min after the injection for determination of plasma [^14^C]2DG radioactivity. At the end of the experiment, animals were anesthetized with an intravenous injection of sodium pentobarbital and tissues were collected and immediately frozen in liquid nitrogen for later analysis of tissue [1-^14^C]-2-deoxyglucose phosphate ([^14^C]2DGP) radioactivity. Blood glucose was measured using an Accu-Chek glucometer (Roche, Germany). Plasma insulin was measured using the Linco rat/mouse insulin ELISA kits. For determination of plasma radioactivity of [3-^3^H]glucose and [1-^14^C]2DG, plasma samples were deproteinized with ZnSO_4_ and Ba(OH)_2_ and counted using a Liquid Scintillation Counter (Beckman Coulter LS6500 Multi-purpose Scintillation Counter). Glucose turnover rate, hepatic glucose production and tissue glucose uptake were calculated as described elsewhere [40,41,43].

### Insulin Tolerance Test

Insulin tolerance tests were performed in high fat fed mice at 24 weeks of age. The day prior to the test, body composition was determined by magnetic resonance (Echo MRI1100, Houston, TX) and lean mass values were used to calculate insulin dose (1 U/kg of lean mass; Humulin R-100, Lilly, U.S.A). On the day of the test, fasting blood glucose concentrations were determined following a 6 hr fast, after which mice received an intraperitoneal injection of insulin diluted in sterile PBS. Blood glucose concentrations were measured every 15 min over a two-hour period post-injection (One Touch Ultra2 hand-held glucometer, LifeScan Europe, Zug, Switzerland).

### RNA Sequencing Analysis and Bioinformatics

Total RNA was extracted from *m. quadriceps femoris* using a Pure Link RNA mini kit (Life Technologies) and then analyzed using an Agilent Bioanalyzer DNA High Sensitivity kit. All samples had a RNA Integrity numbers >7.9. The RNA (1 μg) was enriched for Poly A RNA using an Ambion Dynabeads mRNA Direct Micro kit and barcoded libraries for sequencing were prepared using the Life Technologies RNAseq V2 kit for Ion Torrent according to manufacturer’s standard protocol. The libraries were pooled based on the concentration of each sample between 200-350bp, purified on a Pippin Prep gel, quantified by the Agilent Bioanalyzer and sequenced on an Ion Torrent Proton sequencer. Alignments were made to the mouse genome GRCm38.75 using Tophat 2.0.10 [44] and Bowtie 1.0.0 [45] to incorporate color space data. Counts tables were generated using HTSeq version 0.5.4p5 [46]. Differential expression analyses were performed using DESeq2 version 1.20.0 [47]. All results are presented in Supplementary Table 1, and deposited into the Gene Expression Omnibus as GSE84312. To compare our results to other gene-sets we performed Gene Set Enrichment Analyses (GSEA) comparing our rank-ordered gene lists to annotated gene sets from Gene Ontology, KEGG, Biocarta, Reactome, TRANSFAC and CGP provided as part of MSigDB v6.2 [48,49]. All pathways that met significance at an adjusted p-value of 0.25 are presented in Supplementary Table 2. For comparison of differentially expressed genes, we re-analyzed the *Tsc2* knockout MEFs from GSE21755 [50] and compared with our differentially regulated gene sets.

### Western Blotting

Protein lysates were generated from *m. quadriceps femoris* in RIPA buffer (50 mM Tris pH 7.4, 0.25% sodium deoxycholate, 1% NP40, 150 mM sodium choride, 1 mM EDTA, 100 μM sodium vanadate, 5mM sodium fluoride, 10 mM sodium pyrophosphate and 1X protease inhibitors) or HTNG buffer (50 mM HEPES, pH 7.4, 150 mM sodium chloride, 10% glycerol, 10% triton X-100 and 1X protease inhibitors) by mechanical disruption in a Qialyser for 5 minutes at 30Hz. Lysates were clarified at 14 000 RPM for 15 minutes and quantified by Bradford assays. Proteins were separated by SDS-PAGE, transferred to nitrocellulose and blotted with antibodies described in the figure legends. Primary antibodies used in this study were raised against pS6 (pSer236/236, Cell Signaling #2211), S6 (Cell Signaling #2317), GAPDH (Proteintech #10494) and Sarcolipin (EMD Millipore #ABT-13). Near infra-red secondary antibodies raised against rabbit (Alexa Fluor #A21109) and mouse (Alexa Fluor 790 #A11371) were used to visualize blots on a LiCor Odyssey. Relative protein abundance was quantified using Image Studio Lite software.

### NADH Tetrazolium Reductase Staining

For histology, muscles (*m. quadriceps femoris*) were frozen in liquid nitrogen-cooled isopentane, mounted in OTC and sectioned using a cryostat to 10 µm thickness. Frozen sections were incubated at 37°C for 30 min in pre-warmed 200 mM Tris buffer pH7.4, containing 245 µM nitro blue tetrazolium and 1.13 mM NADH. Sections were rinsed in water, dehydrated, and mounted under coverslips. Staining was visualized and photographed using an Evos XL Core transmitted-light inverted imaging system (Thermo Fisher Scientific).

### Statistical Analyses

All statistical analyses were performed using R, version 3.2.2 [51]. For longitudinal measurements (body weights, fat mass and lean mass), data were analyzed by mixed linear models using uncorrelated random slopes and intercepts using the lme4 package version 1.1-8 [52]. Statistical significance was determined via likelihood ratio tests between models containing or missing the genotype term. Pairwise comparisons were tested for normality via a Shapiro-Wilk test, and for equal variance via Levene’s test. For survival analyses and Cox proportional hazard tests, the survival package was used (version 2.38-3, [53,54]). We tested the assumptions of proportional hazards (with Shoenfeld residuals) and found no significant deviation from this assumption (p=0.875). Based on these, appropriate pairwise tests were performed as indicated in the figure legends. Corrections for testing of multiple hypotheses were done using the method of Benjamini and Hochberg [55]. Statistical significance was designated at p/q<0.05 for all assays, except GSEA analyses where q<0.25 was used. All raw data and statistical analyses for this manuscript are available at http://bridgeslab.github.io/TissueSpecificTscKnockouts.

## Results

### Rapamycin Treatment Blunts High Fat Diet-Induced Increases in Energy Expenditure

To test whether mTORC1 plays a role in the short-term thermogenic responses to obesogenic diets, we measured the total energy expenditure of C57BL6/J mice during a dietary shift between low fat (chow) and HFD in the presence or absence of the specific mTOR inhibitor rapamycin. As depicted in Figure 1A, individually housed animals were vehicle-injected daily for four days, followed by three days of either vehicle or rapamycin injection. After three days of treatment, all animals were moved from a chow diet to HFD. As shown in Figure 1B, the switch to HFD caused an 8.1% increase in total energy expenditure in the vehicle injected mice during the dark phase and a 6.4% increase during the light phase. Compared to vehicle treated mice, rapamycin injection suppressed the HFD-induced increase in energy expenditure (p=2.1× 10^−4^). Notably, these effects were not associated with decreases in physical activity (Figure 1C). These data support the hypothesis that mTORC1 is required for the increase in energy expenditure observed in response to acute HFD feeding.

**Figure 1:**
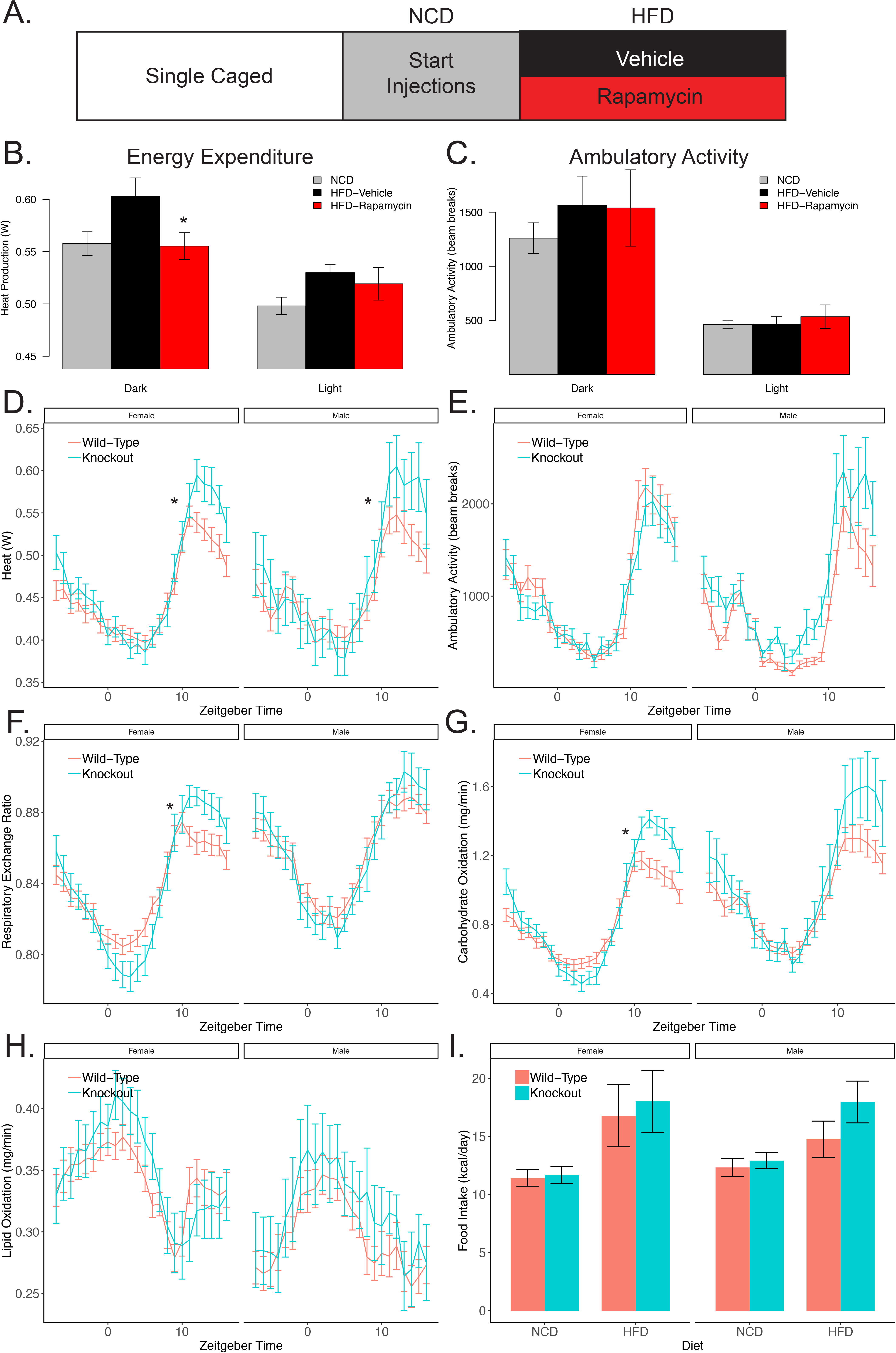
mTORC1 regulates energy expenditure. A) Summary of the rapamycin/high fat diet experimental protocol, and its effect on B) energy expenditure and C) ambulatory activity of 10-week-old male C57BL/6J mice. D) Total energy expenditure, E) ambulatory activity, F) respiratory exchange ratios (RER) and G) rates of carbohydrate oxidation and H) fat oxidation of 10-week-old mice with *Ckmm-Cre* driven knockout of *Tsc1* and their control littermates. H) Absolute food intake of muscle *Tsc1-*knockout mice and their control littermates that received either normal chow diet (NCD) or a high fat diet (HFD). Food intake data were determined over 4-12 weeks on a per-cage basis for mice housed by genotype (n=5-20/group).

### Activation of mTORC1 in Muscle Increases Energy Expenditure

To test whether skeletal muscle mTORC1 activation would result in increases in energy expenditure, we performed indirect calorimetry studies on *Ckmm-Cre* driven *Tsc1* knockout mice. We observed increased total energy expenditure in muscle specific *Tsc1* knockout mice (Figure 1D, p<1 × 10^−6^), and the magnitude of this difference was greater during the dark (active) phase (7.0% increase in males, 6.8% increase in females). Although the greatest energy expenditure differences primarily occurred during the active phase, we did not observe any differences in physical activity (Figure 1E). While there were no significant differences in the respiratory exchange ratio (RER) between knockout and control male mice (Figure 1F), female muscle *Tsc1* knockout mice had a lower RER during the dark period (suggestive of greater lipid utilization while active), and a higher RER during the light period (suggestive of greater carbohydrate utilization while resting) compared to their control counterparts. This finding is corroborated by rates of carbohydrate and lipid oxidation (Figure 1 G and H), and may suggest sexual dimorphism in how muscle mTORC1 signaling affects metabolic flexibility. Together, these data are consistent with a physiological role for mTORC1 in moderating organismal energy expenditure.

### Activation of mTORC1 in Muscle does not alter energy intake

We next evaluated the effect of *Ckmm-Cre* driven *Tsc1* knockout on energy intake in animals receiving either standard laboratory chow or HFD. As shown in Figure 1I, mice receiving the HFD ingested more calories than mice receiving chow (p<0.001); however, there were no differences in energy intake between control and muscle *Tsc1* knockout mice within each diet (p=0.426), and no differences between sexes (p=0.785). While not significant, there was a slight elevation in the energy intake of male knockout mice receiving HFD, consistent with previous reports that show mice with *ACTA1-Cre* driven *Tsc1* knockout eat more food relative to their body weight than control mice when provided a diet consisting of 60% calories from fat [5].

### Activation of mTORC1 in Muscle Causes Resistance to Age-and Diet-Induced Obesity

Given our finding that mTORC1 activation in skeletal muscle caused elevated energy expenditure in the absence of increased energy intake or physical activity, we sought to understand the physiological significance of mTORC1 activation on body composition. The body weights (Figure 2A) and composition (Figure 2B-C) of male muscle *Tsc1* knockout mice given a normal chow diet was determined weekly, over the course of 7 months. As animals aged, control mice accreted body fat, whereas fat mass gains were minimal in the knockout group; a striking 84% difference being observed between knockout and control groups (Figures 2B, p=1.7 × 10^−10^). Previous work using *ACTA1-Cre* mediated knockout of *Tsc1* also report lower body fat accumulation in knockouts compared to control mice [5,6], concomitant with reductions in lean mass [6,21], the latter finding not replicated in this study (Figure 2C, p=0.743 at endpoint). To determine if reductions in body fat gains were adipose depot-specific, we determined the weights of subcutaneous (dorsolumbar-inguinal) and visceral (epididymal) fat pads from male control and *Tsc1* knockout mice, and found that both fat depots were markedly lighter (decreased 79% and 76%, respectively, each p<0.0001; Figure 2D). Together, these findings demonstrate that skeletal muscle mTORC1 activation results in lower adiposity gains across the lifespan.

**Figure 2:**
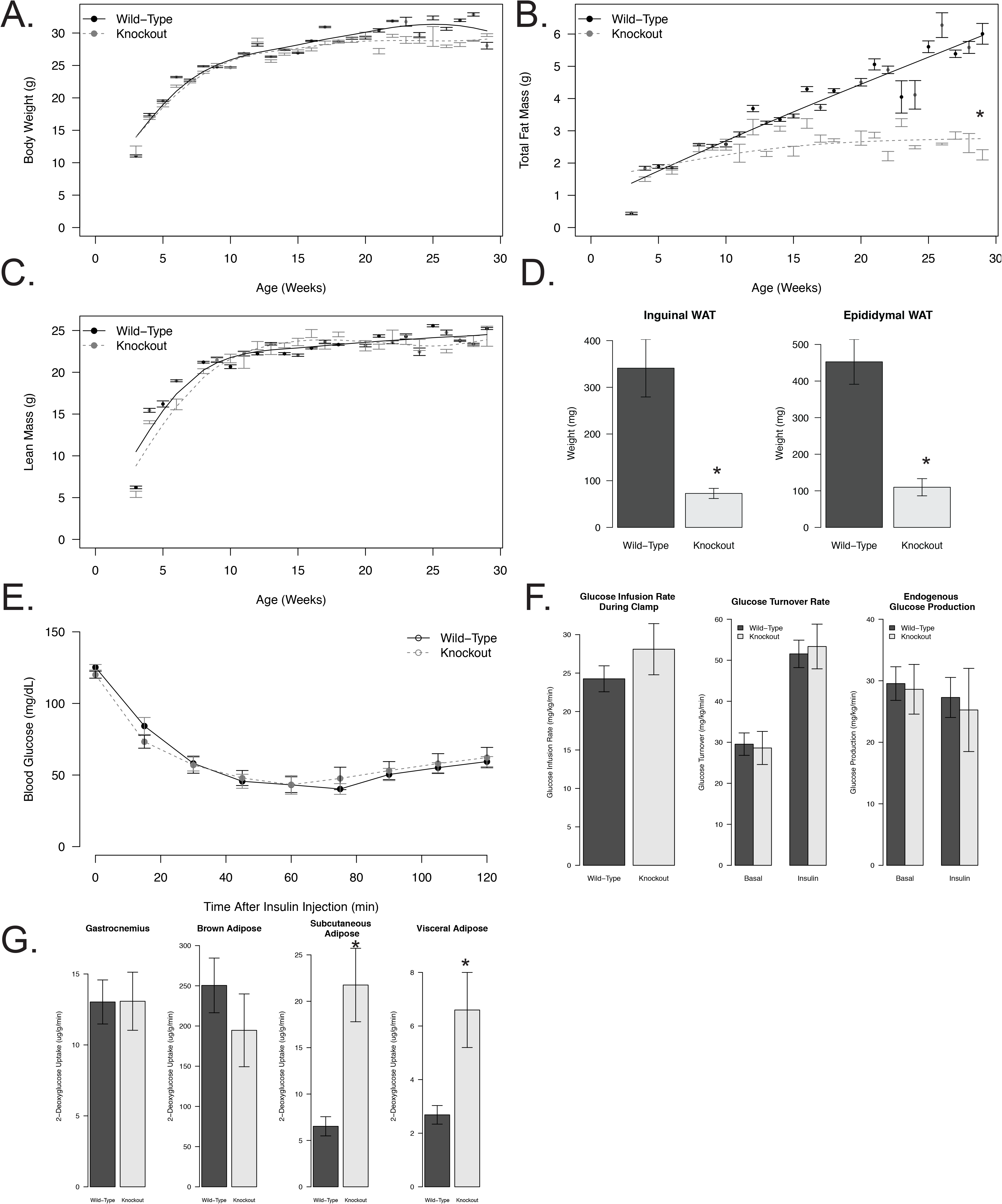
Skeletal muscle mTORC1 activation attenuates age-associated gains in adiposity. A) Body Weight, B) Total body fat and C) lean mass of male mice with *Ckmm-Cre* driven knockout of *Tsc1* and their control littermates, as determined longitudinally, from the age of weaning until 29-weeks of age. D) Weights of the dorsolumbar-inguinal and gonadal fat depots. E) Blood glucose curves during an tolerance test and F) the glucose infusion, glucose turnover and endogenous glucose production rates during a hyperinsulinemic euglycemic clamp. G) Tissue 2-deoxyglucose uptake under insulin stimulated conditions at the end of the clamp. Statistical significance is denoted by asterisks indicating p<0.05, based on a likelihood ratio test (B) or Mann-Whitney test (D and G, due to lack of normality). Data are reported for n=7 muscle *Tsc1* knockout mice and n=25 control mice. Mice were fasted 5-6 hours prior to insulin tolerance tests and hyperinsulinemic clamp studies.

### Activation of mTORC1 in Muscle Does Not Induce Systemic Insulin Resistance

Activation of mTORC1 has been reported to induce systemic insulin resistance in some systems, so we next determined insulin sensitivity in these mice by insulin tolerance test. As shown in Figure 2E, male knockout mice had similar insulin responsiveness to control mice. To pursue this unexpected finding further, we performed hyperinsulinemic euglycemic clamps, and found similar results. The rate of glucose infusion during the clamp was not different in knockout mice, nor were glucose turnover rates (Figure 2F) or the accumulation of 2-deoxyglucose in gastrocnemius muscles (Figure 2G). 2-deoxyglucose uptake into brown adipose tissue was not different between knockout and control mice, whereas 2-deoxyglucose uptake into both subcutaneous and visceral white adipose tissue depots was markedly elevated.

To determine if a palatable, hypercaloric diet would induce changes in body composition or insulin sensitivity in mice with *Ckmm-Cre* driven knockout of *Tsc1*, we placed mice on a diet containing 45% of calories from fat and found that both male and female mice were resistant to HFD-induced weight gain. (Figure 3A) Differences in body weight were primarily due to differences in fat mass, which, compared to control mice, was 60% lower in knockout males and 58% lower in knockout females by the end of the study (Figure 3B, p<1.0×10^−6^ for each). These data are consistent with previous reports in *ACTA1-Cre* mediated *Tsc1* knockout mice fed a diet containing 60% calories from fat [5,6]; however, in our model lean masses across both sexes were not different between control and knockout mice on HFD (Figure 3C, p=0.941). Consistent with the *in vivo* body composition data, we observed a 75-80% difference in the weights of both the gonadal and inguinal fat pads from male and female knockout mice compared to their relative control groups (all p<0.001; Figure 3D). Together, these findings demonstrate that mTORC1 activation in skeletal muscle can protect against adiposity gains under otherwise obesogenic conditions.

**Figure 3:**
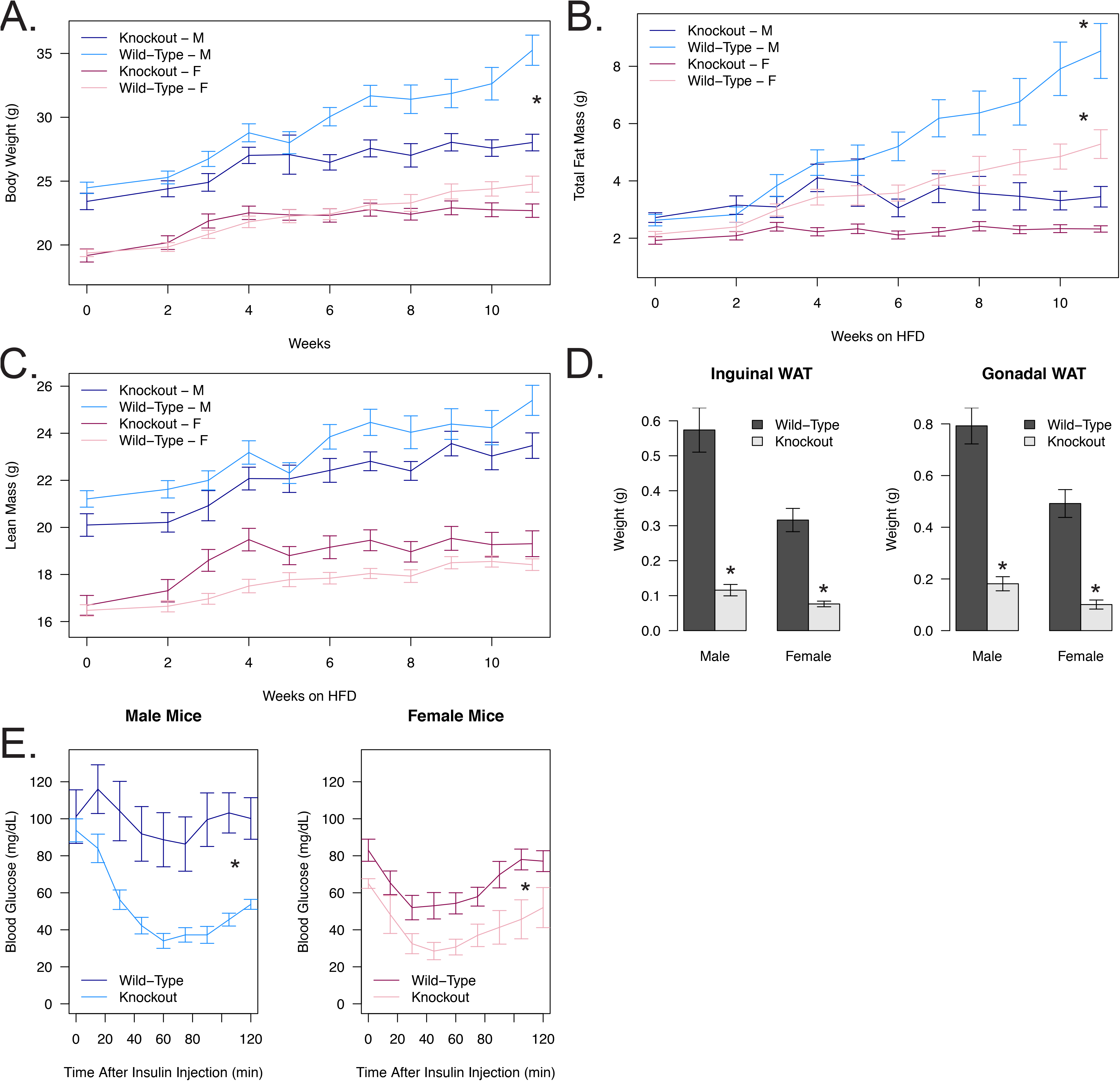
Skeletal muscle mTORC1 activation protects against diet-induced obesity and insulin resistance. A) Total body weight, B) total body fat and C) total lean mass of male and female mice with *Ckmm-Cre* driven knockout of *Tsc1* and their control littermates following 11-weeks of an obesogenic diet containing 45% energy from fat (HFD), beginning at 10-weeks of age. D) Weights of the dorsolumbar-inguinal and gonadal fat depots after 11-weeks of HFD. E) Blood glucose concentration curves during an insulin tolerance test in 6 hour-fasted male and female mice with *Ckmm-Cre* driven knockout of *Tsc1* and their control littermates after 11-weeks of HFD. Statistical significance (*p<0.05, n=5/7) was determined via a Welch’s *t* test (D, males), or a Mann-Whitney test (D, females, due to lack of normality).

As shown in Figure 3E, compared to control mice, both male and female muscle *Tsc1* knockout mice were more insulin responsive (33% reduction in the area under the blood glucose curve for females, 45% difference for male mice; p=0.045 and 0.014 respectively). This finding is consistent with the hypothesis that muscle *Tsc1* knockout mice are not lipodystrophic but have improved glycemic control in addition to reduced adiposity. We propose that mice with muscle *Tsc1* knockout are protected from HFD-induced adipose tissue expansion as a result of having chronically elevated energy expenditure, and that the attenuated body fat gains observed in muscle *Tsc1* knockout mice facilitates the preservation of insulin sensitivity in the face of a HFD.

### Muscle mTORC1 Activation Causes Enrichment of Gene Sets Involved in Fatty Acid Uptake and Amino Acid Uptake

To gain further insight into the mTORC1 activity-driven mechanisms within skeletal muscle that increase energy expenditure and limit body fat accumulation in *Tsc1* knockout mice, we performed RNA sequencing studies in RNA obtained from *m. quadriceps femoris* from chow-fed male mice with *Ckmm-Cre* driven *Tsc1* knockout and their floxed litter mates. We identified 4403 differentially expressed gene transcripts in these animals, including 2464 upregulated genes and 1939 downregulated genes (see Supplementary Table 1 for complete list). To identify the pathways and networks associated with these differentially expressed gene transcripts, we performed gene-set enrichment analyses, finding 674 differentially regulated gene sets (see Supplementary Table 2). Among the significantly enriched gene sets were genes also regulated by *Tsc2* deletion in MEFs, and by treatment with rapamycin [50,56], indicating there are a core set of mTORC1 dependent genes that are similarly regulated across different tissues. Consistent with this observation, we found that 58% of the differentially expressed genes in the *Tsc1* knockout muscles overlapped with previously published differentially expressed genes in *Tsc2* knockout MEFs [50]. Other gene sets we identified as being upregulated by *Tsc1* ablation in skeletal muscle include IGF1 targets in MCF-7 cells [57], genes involved in protein synthesis, amino acid (Figure 4A) and fatty acid uptake (Figure 4B), and calcium trafficking (Figure 4C),. Most amino acid transporters were increased at the mRNA level (Figure 4A), while the fatty acid binding protein *Fabp3* was also increased at the transcriptional level (Figure 4B).

**Figure 4:**
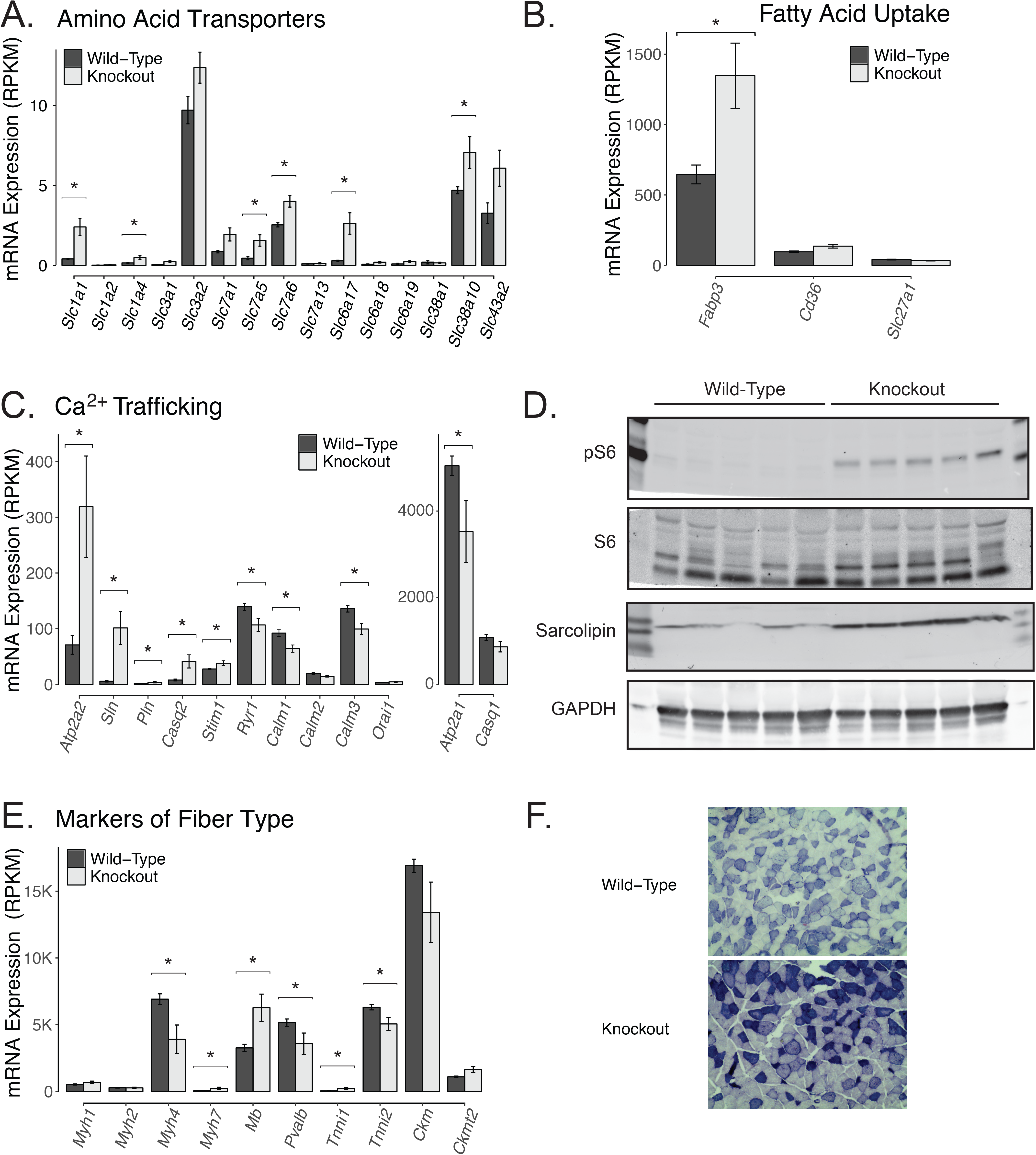
Skeletal muscle mTORC1 activation alters the transcriptional regulation of nutrient uptake and oxidative capacity. A) Expression of amino acid transporters B) fatty acid transporters, C) Ca^2+^ trafficking and, markers of fiber type E) as determined by RNAseq. D) Representative protein expression of S6 phosphorylation at Ser236/236, total S6 and sarcolipin in quadriceps muscles from male control mice and mice with *Ckmm-Cre* driven knockout of *Tsc1*. F) Representative images of quadriceps muscle from mice with *Ckmm-Cre* driven knockout of *Tsc1* and their control littermates stained for NADH-tetrazolium reductase, where oxidative fibers stain darkest. Asterisks indicates adjusted p-values of <0.05.

### Muscle mTORC1 Activation Increases Thermogenic Signaling via Alterations in Intramyocellular Ca^2+^ Dynamics

To identify the molecular mechanisms causing increased energy expenditure in muscle *Tsc1* knockout mice, we determined the expression of transcripts known to be important contributors to skeletal muscle thermogenesis. We observed dramatic increases in the ATP-dependent SR/ER Ca2+ pump SERCA2 (encoded by *Atp2a2*, see Figure 4C), and its un-coupler Sarcolipin (encoded by *Sln*; Figure 4C), proteins previously reported as playing an integral role in muscle-specific thermogenesis [58–61]. At the protein level, Sarcolipin was increased 4.1-fold (p=4.5 × 10^−6^; Figure 4D, pS6 is shown as a positive control for mTORC1 activation). We propose that the increased energy expenditure observed in mice with muscle-specific *Tsc1* ablation may be caused, in part, by increased futile cycling of Ca^2+^ by uncoupled SERCA2 (therefore increasing ATP hydrolysis) at the SR. Consistent with this hypothesis, we observed increases in the expression of other transcripts important for Ca^2+^ trafficking, including *Pln*, *Casq2, Stim1* (Figure 4C), *Mfn1-2* (Supplementary Table 1) and the subunits of the mitochondrial calcium importer (*Mcu, Micu1* and *Micu2*; Supplementary Table 1). We also observed reductions in *Ryr1, Calm1* and *Calm3* expression (Figure 4C), and reductions several plasma membrane Ca^2+^ transporters (see Supplementary Table 1), changes that are likely adaptive mechanisms to manage increased intracellular Ca^2+^ levels associated with SERCA2 uncoupling.

### Constituent mTORC1 Activation Increases the Oxidative Profile of Skeletal Muscle

We also evaluated transcriptional markers of muscle fiber type and observed increases in markers for more oxidative fiber types, including *Myh7, Mb, Tnnc1, Tnni1* and *Atp2a2*, along with downregulation of markers for glycolytic fibers, including *Myh4, Pvalb, Tnnc2, Tnni2* and *Atp2a1* (Figures 4C and E, and Supplementary Table 1). These data suggest that skeletal muscle mTORC1 activation increases the oxidative profile of skeletal muscle at the transcriptional level. These findings are also supported by our observation that skeletal muscle from *Tsc1* ablated mice has greater NADH-dehydrogenase activity (Figure 4F), and are consistent with findings from previous studies on *ACTA1-Tsc1* knockout muscles that report the accumulation of mitochondrial enzymes and changes in muscle fiber size [20,62].

### Muscle mTORC1 Activation Reduces Lifespan

To determine whether skeletal muscle mTORC1 activation-induced increases in energy expenditure affected lifespan, we monitored muscle *Tsc1* knockout animals without manipulation as they aged. Increased signs of aging, including hunched and scruffy appearances at an earlier age, were observed in knockout mice compared to their control littermates. As shown in Figure 5, muscle-specific *Tsc1* knockout mice died of natural causes earlier than control mice. Based on a Cox-proportional hazard model, the hazard ratio was 4.17-fold higher compared to non-knockout littermates (p=2.0 × 10^−5^).

**Figure 5:**
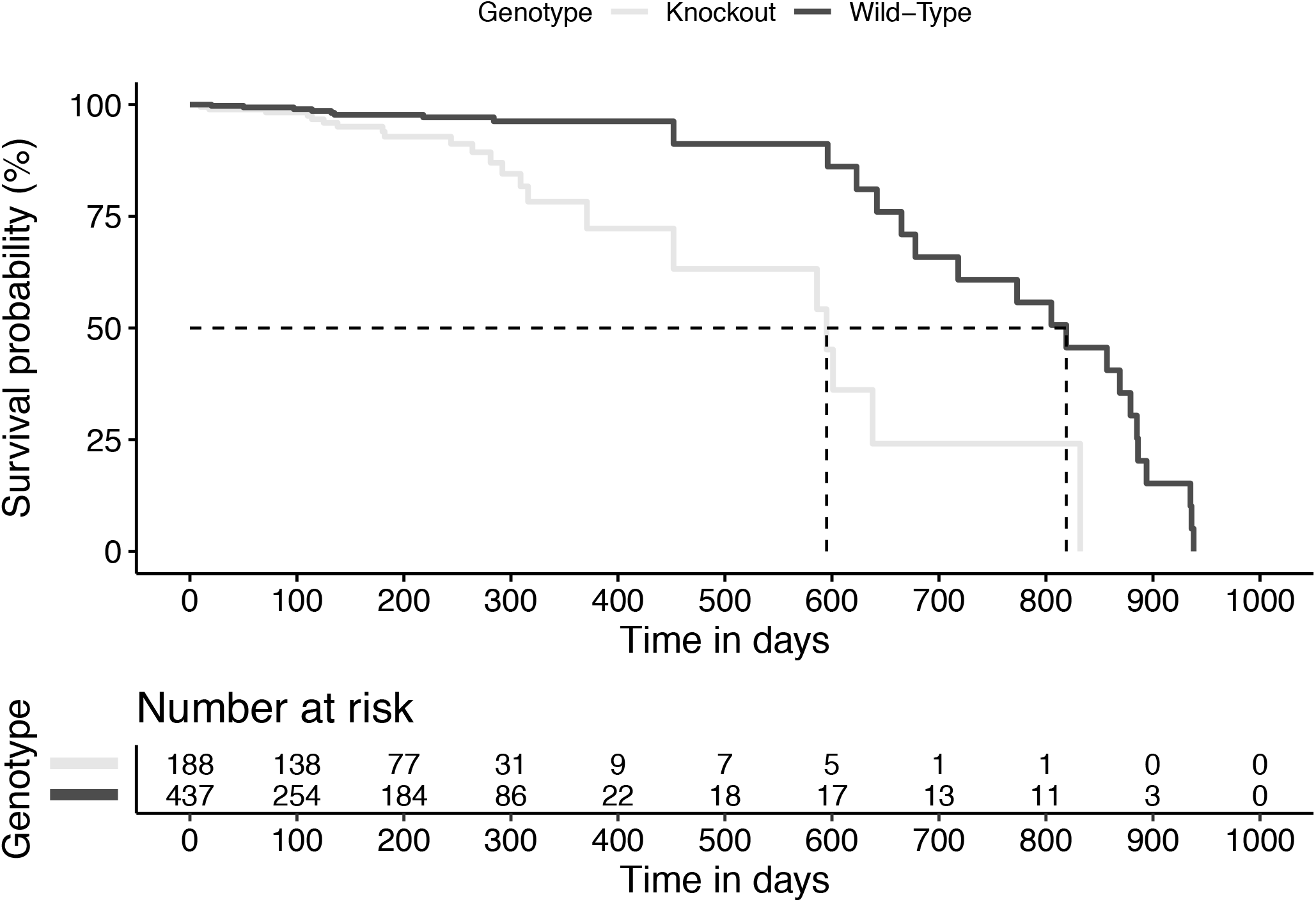
Skeletal muscle mTORC1 activation reduces lifespan in mice. Survival curve of male muscle *Tsc1* knockout mice on a normal chow diet. Dotted lines indicate age at which 50% of animals died.

To determine how muscle *Tsc1* ablation reduces lifespan, a subset of mice were fixed in formalin upon death and sent for veterinary pathology. However, we were unable to identify a consistent cause of death in these mice. In mice with histologic evidence of lesions, the predominant process was neoplasia, and the specific etiology was lymphoma/lymphosarcoma affecting multiple organs, though this was only true for control mice (two out of four) but not knockout animals (none out of three). It is important to note that lack of a specific diagnosis does not necessarily confirm the lack of lesions in examined animals; rather, that autolysis and the small number of animals evaluated may have resulted in loss of identifiable processes or tissues in which an etiology was present in-life.

## Discussion

Here, we show that high fat diet-induced increases in energy expenditure are mTORC1-dependent. We also demonstrate that constitutive activation of skeletal muscle mTORC1 causes elevated total energy expenditure independent of changes in physical activity, and that mTORC1-driven increases in energy expenditure coincide with reduced adiposity, increased insulin responsiveness and the upregulation of skeletal muscle-specific thermogenic mechanisms that involve ATP-dependent futile cycling of Ca^2+^. Consistent with prior work [6,20], we also show transcriptional evidence for an mTORC1-dependent fiber type transition to a more oxidative phenotype, along with other markers of altered substrate oxidation and energy transformation in skeletal muscle.

Skeletal muscle is an important determinant of tissue insulin responsiveness, energy balance and healthy aging. Humans with high baseline grip strength have decreased risk of all-cause mortality [28– 33], whereas interventions that increase muscle mass and strength are associated with improved health outcomes in both young and older populations [63,64]. Given that skeletal muscle is also the primary determinant of resting energy expenditure [65], understanding the molecular mechanisms that influence skeletal muscle health could have important ramifications for the clinical treatment of diseases associated with both obesity and aging.

Our understanding of how mTORC1 regulates skeletal muscle physiology largely consists of the translation-initiation and post-translational role of mTORC1 and its anabolic response to growth factors, nutrients and mechanical loading [9]. Less is known regarding the role of mTORC1 in the regulation of skeletal muscle substrate oxidation and energy expenditure; however, it has been suggested that gene silencing of mTORC1 alters ATP generation through disruption of PGC-1α driven mitochondrial signaling *in vitro* [22], and that the increase in *Ppargc1a* expression that occurs in muscle during the acute post-exercise phase has been shown to be potentiated in mouse skeletal muscle when mTORC1 is inhibited by rapamycin [66]. Previous studies have shown that activating mTORC1 in mice through deletion of *Tsc1* in skeletal muscle results in smaller mice that have significantly lower body fat than control mice and are resistant to both diet-induced obesity and age-associated gains in adiposity [5,6]. Our results agree with these data, and we provide evidence that resistance to the accretion of body fat in these animals may be conferred, in part, by an increase in energy expenditure caused by sarcolipin-driven uncoupling of SERCA2 in skeletal muscle.

Skeletal muscle thermogenic pathways are important contributors to both shivering and non-shivering thermogenesis and while mitochondrial-generated thermogenic pathways have been described as potential targets for leveraging skeletal muscle thermogenesis to combat obesity [67], other heat generating pathways may also be important. The Sarco/Endo-plasmic Reticulum Ca^2+^-ATPase (SERCA) transfers Ca^2+^ from the sarcoplasm into the lumen of the sarcoplasmic reticulum, hydrolyzing ATP in the process. Sarcolipin, a small helical peptide, can interact with SERCA and alter the kinetics of Ca^2+^ re-sequestration by allowing slippage of Ca^2+^ back into the sarcoplasm, thereby ‘decoupling’ Ca^2+^ uptake from SERCA-dependent ATP hydrolysis and creating a futile cycle of Ca^2+^ movement that generates heat [68]. In the present study, *Ckmm-Cre* driven *Tsc1* deletion resulted in increased whole-body energy expenditure (Figure 1D), transcriptional upregulation of both SERCA2 and sarcolipin (Figure 4C), and increased expression of sarcolipin at the protein level (Figure 4D). We predict that the increased amounts of sarcolipin in muscle *Tsc1* knockout mice result in increased SERCA2 uncoupling and higher rates of ATP-dependent futile Ca^2+^ cycling [58,69], a hypothesis supported by our observation that a number of other Ca^2+^ transporters and Ca^2+^ responsive mRNA’s are differentially expressed in the muscles of these mice (Figure 4C and Supplementary Table 1). Increased muscle thermogenesis through the sarcolipin-driven uncoupling of SERCA would increase organismal energy expenditure, thereby resulting in lower adiposity gains. Indeed, this hypothesis is consistent with reports that obesity is exacerbated when *Sln* is ablated [58–60] or prevented when it is *Sln* is overexpressed [61].

In addition to the changes in Ca^2+^ related transcripts, we and others have observed that skeletal muscle-specific activation of mTORC1 via deletion of *Tsc1* results in an increase in the oxidative profile of the skeletal muscle [20]. Furthermore, non-*Tsc1*-driven models of muscle-specific mTORC1 activation, such as those involving knockout of individual components of the GATOR1 complex, result in increased expression of mitochondrial components, particularly TCA cycle intermediates [70], and increased mitochondrial respiration [71]. Conversely, abolishing skeletal muscle mTORC1 activity via Raptor knockout increases mitochondrial coupling efficiency but lowers mitochondrial respiration and reduces the abundance and activities of mitochondrial enzymes [72]. Here, we show that muscle *Tsc1* knockout mice have an increased reliance on carbohydrate oxidation (Figure 1G). Taken together, these observations suggest that mTORC1 influences metabolism by increasing mitochondrial enzyme content, the coupling of oxidative phosphorylation to ATP production, and the dissipation of energy via uncoupling of the sarcoplasmic reticulum.

We cannot rule out other mechanisms linking muscle mTORC1 activity to elevated energy expenditure that may be indirect, such as FGF21 [6,73,74], other myokines or muscle-derived metabolites. It is also worth noting that our observations during the rapamycin experiments are limited in that they do not speak to tissue specificity. Our results are consistent with previous reports demonstrating rapamycin sensitivity in cold-induced thermogenesis [7,8], and while the focus of those studies has been on the important roles of mTORC1 in brown adipose tissue function, they may also speak to the role of mTORC1 in muscle or other thermogenic tissues. Future studies with temporal and tissue-specific loss of mTORC1 function conducted in mice housed at temperatures within their thermoneutral zone will be key to understanding the relative importance of muscle and BAT in both diet- and cold-induced thermogenesis. Furthermore, findings that mTORC1 is important for thermogenesis in both BAT and skeletal muscle may indicate a broader role of mTORC1 in nutrient homeostasis. One response to nutrient overload is to promote anabolism, consistent with mTORC1-dependent activation of protein synthesis, lipogenesis, and glycogenesis [13,35,77]. Thus, it is reasonable to propose that nutrient overload may promote ineffective catabolism as a way of reducing systemic nutrient stress.

Transcriptional profiling across species has identified downregulation of mitochondrial genes in skeletal muscle as a common aging signature [78,79], whereas loss of skeletal muscle mitochondrial function is associated with age-related sarcopenia in *C. elegans* [80,81] and mice. Candidate gene studies on aging have implicated genes with important roles in skeletal muscle metabolism, including *IGF1R*, *AKT1* and *FOXO3A* [82,83], genes that are also linked to mTORC1 signaling. In humans, polymorphisms in *FOXO3A* have been associated with lengthened lifespan [83–89], whereas both mouse and fruit fly models of *FOXO3A* loss of function result in stronger and longer living model organisms [90–92]. Indeed, nonagenarians show downregulation of mTOR pathway genes [93], supporting a role for decreased mTOR signaling in human longevity, whereas in rats, inhibition of mTORC1 via rapalog treatment ameliorates age-related sarcopenia [94]. Here, we show that despite an apparent increase in the oxidative phenotype of muscle, constituent activation of mTORC1 in skeletal muscle decreases lifespan in mice, a finding in consensus with other models of mTORC1 activation [25–27].

## Conclusions

We have shown that increases in energy expenditure following a high fat diet-are mTORC1-dependent and that elevated energy expenditure caused by ablation of *Tsc1*, and thus constituent activation of skeletal muscle mTORC1, coincides with the upregulation of skeletal muscle-specific thermogenic mechanisms that involve the sarcolipin-driven futile cycling of Ca^2+^ through SERCA2. These findings support the hypothesis that activation of mTORC1 and its downstream targets, specifically in skeletal muscle, may play a role in adaptive thermogenesis, and point to a role for mTORC1 in stimulating mechanisms of energy expenditure in response to caloric overload. Whether mTORC1-dependent Ca^2+^ cycling in muscle is an effective therapeutic strategy for targeting weight loss remains to be determined, but given the negative effects of mTORC1 activation on lifespan, clinical utility is expected to be low. Future studies will confirm whether mTORC1 exerts its effects on skeletal muscle thermogenic pathways directly or indirectly and will identify whether the positive effects of skeletal muscle mTORC1 activation can be separated from the negative effects to reduce adiposity and protect against obesity.

## Acknowledgements

The authors would like to thank Melanie Schmitt of the UM Animal Phenotyping Core for assistance with CLAMS and glucose clamp studies on the muscle *Tsc1* knockout mice. William Taylor, Caitlin Costelle and Felicia Waller at the UTHSC Molecular Resource Center provided support for the transcriptomic studies. We would also like to thank the other members of the Bridges, Han, and Saltiel laboratories for helpful discussions regarding this project.

This work was supported by Le Bonheur Grant 650700 (DB), NIH Grants DK107535 (DB), DK076906 and DK117551 (ARS), funds from the Memphis Research Consortium (DB and JCH), the Center for Integrative and Translational Genetics (DB), the UTHSC Department of Physiology Qiugley Award (IH) and a Rackham Merit Fellowship (MCM). This work also utilized Core Services supported by NIH grants DK089503, DK110768, DK020572 and AR069620 to the University of Michigan.

## Supplementary Table Legends

**Supplementary Table 1: Gene expression differences in muscle *Tsc1* knockout quadriceps.** Full results of differential expression analysis.

**Supplementary Table 2: Gene set enrichment analysis of *Tsc1* knockout quadriceps.** All pathways that met an adjusted p-value of 0.25 are shown. NES (normalized enrichment score) indicates pathway effect size with positive numbers indicating positive enrichment of this gene set in these data. Gene details are the genes which drove this positive or negative association. Both nominal (NOM) and FDR adjusted (FDR) p/q values are shown.

